# On the emergence of structural complexity in RNA replicators

**DOI:** 10.1101/218990

**Authors:** Carlos G. Oliver, Vladimir Reinharz, Jérôme Waldispühl

## Abstract

The RNA world hypothesis relies on the ability of ribonucleic acids to spontaneously acquire complex structures capable of supporting essential biological functions. Multiple sophisticated evolutionary models have been proposed for their emergence, but they often assume specific conditions. In this work we explore a simple and parsimonious scenario describing the emergence of complex molecular structures at the early stages of life. We show that at specific GC-content regimes, an undirected replication model is sufficient to explain the apparition of multi-branched RNA secondary structures – a structural signature of many essential ribozymes. We ran a large scale computational study to map energetically stable structures on complete mutational networks of 50-nucleotide-long RNA sequences. Our results reveal that the sequence landscape with stable structures is enriched with multi-branched structures at a length scale coinciding with the appearance of complex structures in RNA databases. A random replication mechanism preserving a 50% GC-content may suffice to explain a natural enrichment of stable complex structures in populations of functional RNAs. By contrast, an evolutionary mechanism eliciting the most stable folds at each generation appears to help reaching multi-branched structures at highest GC content.

## 1 Introduction

RNA are versatile molecules that can virtually fulfill all fundamental needs and functions of the living, from storing information to catalyzing chemical reactions and regulating gene expression. The RNA world hypothesis [12, 18] builds upon this observation to describe a scenario of the emergence of life based on RNAs. Despite criticisms [37, 47, 56], recent studies presented plausible paths towards an early assembly of RNA molecules has contributed to strengthening this hypothesis [6, 24, 40, 42, 46]. Yet, the emergence of nucleic acids is only one part of this problem. We also need to elucidate the mechanisms that enabled the discovery of functional molecules and the transmission of genetic information [64].

Many non-coding RNAs acquire functions through structures. Classically, we describe these structures at two levels of abstraction. The secondary structure encompasses the Watson-Crick and Wobble base pairs, while the tertiary structure describes the 3D coordinates of all atoms. Noticeably, RNA secondary structures are more conserved than sequences and thus provide a reliable signature of RNA function [34]. Moreover, many essential molecular functions are supported by nucleic acids with complex shapes often characterized by the occurrence of a k-way junction (a loop connecting three or more stem-like regions also known as multi-loop; See Fig. 1C) in their secondary structure (e.g. 5s rRNA, tRNA, hammerhead ribozyme). A theory describing how RNA populations evolve to “discover” these functional multi-branched secondary structures is therefore an important step toward a validation of the RNA world hypothesis [22].

**Figure 1:**
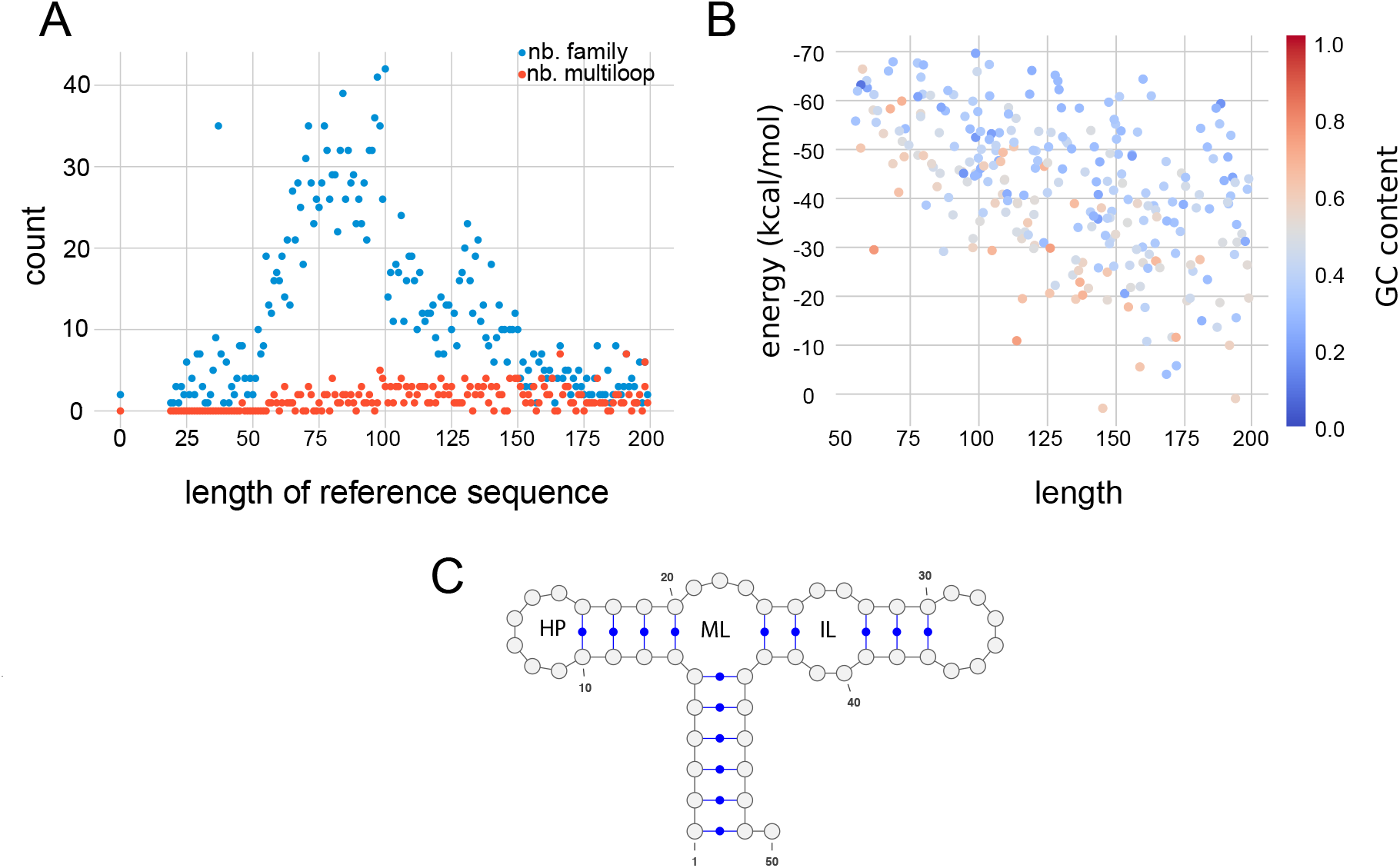
Statistics on Rfam families [34]. (A) We plot the number of families with respect to the average length of the sequences in these families. Red dots show the numbers of families with a consensus structure that contains a multi-loop, while blue dots show those without. (B) We plot the average folding energy and length of sequences for each Rfam families having multi-branched consensus structures. The color indicates the average GC content of the family. (C) An illustration of an RNA 2D fold (generated using VARNA [10]) and label its secondary structure elements as internal loop (IL), multi-loop (ML) and hairpin (HP).

Since the first analysis of RNA neutral networks (i.e. networks of RNA sequences with identical structures) [45], computer simulations are the method of choice for characterizing the evolutionary landscape and population dynamics of structured RNA molecules. Indeed, secondary structures can be reliably predicted from sequence data only [31], allowing fast and accurate prediction of phenotypes (i.e. secondary structure) from genotypes (i.e. sequence).

A large body of literature has been dedicated to the computational analysis of RNA sequence-structure maps (i.e. genotype-phenotype maps) and properties of RNA populations evolving under natural selection. In a seminal series of papers P. Schuster and co-workers set up the basis of a theoretical framework to study the evolutionary landscape of RNA molecules, and used it to reveal intricate properties of networks of sequences with the same structure (a.k.a. neutral networks) [14, 20, 45, 51, 53, 55]. This work inspired numerous computational studies that refined our understanding of neutral models [2, 3, 11, 69, 73], as well as kinetics of populations of evolving nucleic acids [27, 59, 61].

In this paper, we perform computer simulations to study the evolutionary mechanisms that enabled the emergence of multi-branched secondary structures. This feature turns out to be relatively common even for short RNAs. An analysis of the consensus secondary structures available in the Rfam database reveals that multi-loops can be found in approximately 10% of RNA families whose average size of sequences ranges from 50 to 100 nucleotides (See Fig. 1A). This observation contrasts with earlier studies that revealed that the vast majority of predicted minimum free energy (MFE) secondary structures obtained from a uniform sampling of shorter RNA sequences (i.e. 35 nt.) are stem-like structures (i.e. no multi-loop) [60], which render a spontaneous emergence of complex structures (i.e. secondary structures with a multi-loop) unlikely on such short sequences. Nonetheless, computational studies of RNA sequence-structure maps showed that neutral networks percolate the whole sequence landscape [16, 54]. Even though this property undeniably augments the accessibility of target structures, the size and connectivity of neutral networks also decreases drastically with the complexity of structures.

In the most commonly accepted scenarios, the establishment of a stable, autonomous, and functional self-reproductive molecular system subject to natural selection, relies on the presence of polymerases [22]. Such molecules are long (200 nt.) and thus unlikely to be discovered randomly. Instead, it has been suggested that evolution proceeded by stages [30]. Polymerases were assembled from smaller monomers (*∼*50 nt.) that are more likely to emerge from prebiotic chemistry [21, 68]. At this point, and not before, parallel natural selection processes of specific functional structures could be triggered.

Interestingly, *in vitro* experiments revealed the extreme versatility of random nucleic acids [4, 5, 43, 50], and suggested that essential RNA molecules such as the hammerhead ribozyme could have multiple origins [49]. All together, these observations reinforce the plausibility of a spontaneous emergence of multiple functional sub-units. But they also question us about the likelihood of such events and the existence of intrinsic forces promoting these phenomena.

Various theoretical models have been proposed to highlight mechanisms that may have favoured the birth and growth of structural complexity from replications of small monomers. Computational studies have been of tremendous help to explore various scenarios. In particular, numerical simulations enabled us to study the effects of polymerization on mineral surfaces [7, 62] or the importance of spatial diffusion [57]. Noticeably, the majority of these scenarios are proposing a development of RNA structural complexity outside a cellular barrier. But this assumption results in major challenges. The first of them is to explain how a system that evolved and adapted in an exposed milieu would transition to a membrane-protected environment. In particular, the presence of a membrane would radically change the tradeoff between stability and complexity of RNA structures. Indeed, stable folds often lack the complexity necessary to support novel functions but are more resilient to harsh pre-cellular environments [25].

In this work, we aim to characterize the structural repertoire accessible by replicating RNA populations. This scenario is compatible with the hypothesis of an early development of membranes [1, 8, 35, 64] and does not require invoking hybridization on surfaces, or the presence of large self-replicating ribozymes [39]. Importantly, we exclude directed evolution scenarios characterized by a progressive adaptations to a phenotype (e.g replication with errors minimizing the distance of Minimum Free Energy (MFE) structures to a target structure [52]). Instead, we rely on intrinsic forces (i.e. increasing secondary structure stability) for driving the evolutionary process and study the impact of GC-content bias.

We apply customized algorithms to study the distribution of structures accessible to all mutant sequences with 50 nucleotides [71, 72]. This approach considerably expands the scope and significance of comprehensive RNA evolutionary studies that were previously limited to sequences with less than 20 nucleotides [9, 20], or restricted to explore a small fraction of the sequence landscape of sequences [11, 60]. Our simulations reveal that based on the strength of the selective pressure applied on the replicating population, different GC content biases facilitate the emergence of complex secondary structures with k-way junctions. In the absence of selective pressure, low to medium GC contents (0.3-0.5) may suffice to explain the distribution of multi-loop structures observed in databases. By contrast, high GC contents help populations eliciting sequences with stable folds to discover structure with k-way junctions. These results provide valuable insights into previous contributions studying GC content biases in stability-flexibility tradeoffs [28], prebiotic nucleotide distributions [17, 41] and the accessibility of complex phenotypes [60].

## 2 Results

### 2.1 Our approach

We apply complementary techniques to explore the RNA mutation landscape and characterize the structures accessible from an initial pool of random sequences under distinct evolutionary scenarios (See Fig. 2). Importantly, our analysis explicitly models GC content bias to understand the effect of potential nucleotide scarcity in prebiotic conditions [17, 41].

**Figure 2:**
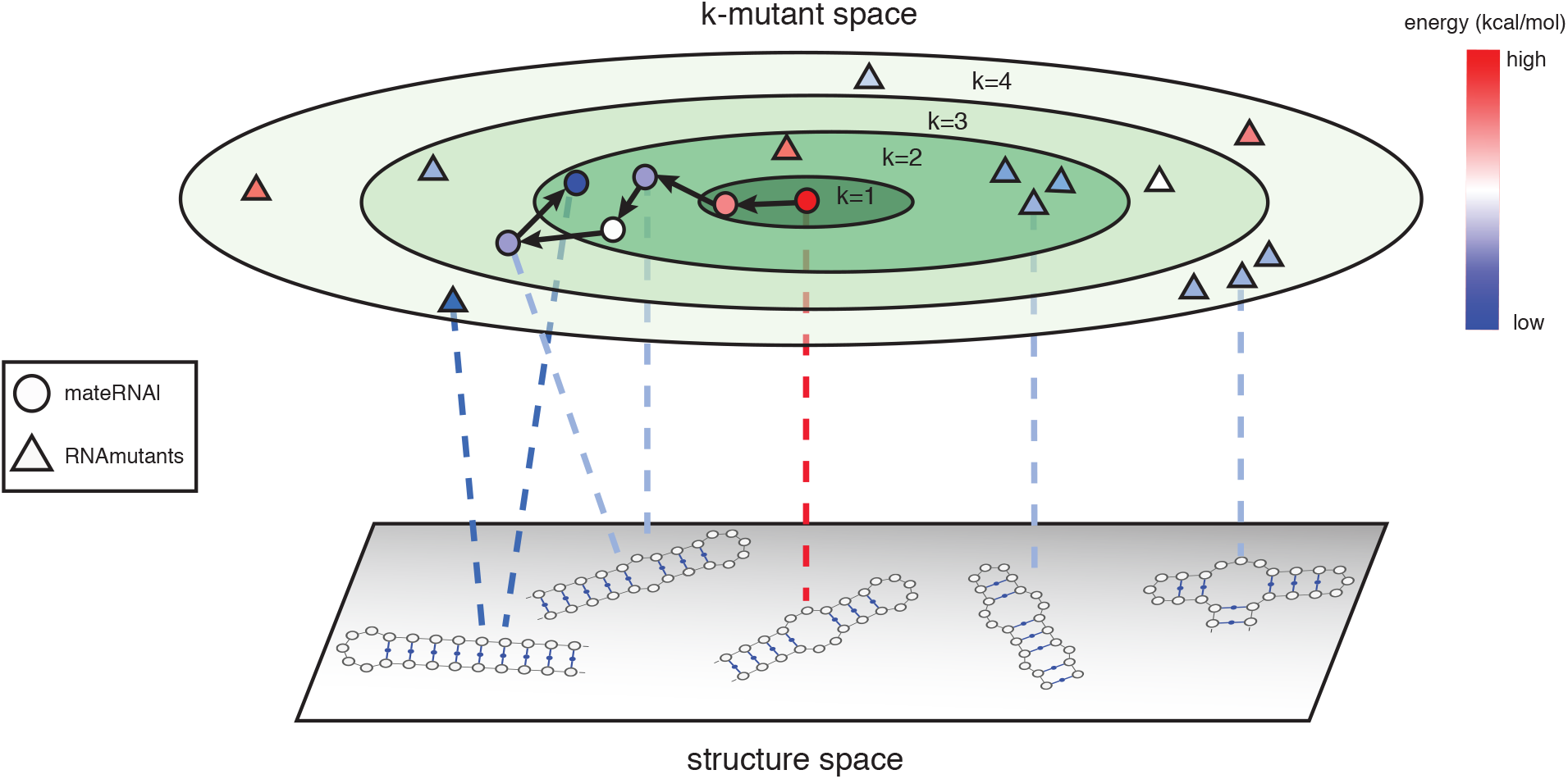
*Illustration of mutational sampling methods.* Concentric ellipses represent the space of sequence k-mutant neighbourhoods around a root sequence pictured at the centre of the ellipses. Each ring holds all sequences that are *k* mutations from the root. We show the contrast between an evolutionary trajectory (`mateRNAl`) along this space and mutational ensemble sampling (`RNAmutants`) represented respectively as circles and triangles. The layer below the mutational space represents the space of all possible secondary structures, and dotted lines illustrate the mapping from sequences to structures. The colour of the sampled mutants denotes the energy of the sequence-structure pair sampled. In both sampling methods, sequence-structure pairs with lower energies are favoured. Evolutionary sampling is always limited to explore sequences accessible from the parental sequence and so we have arrows pointing from parents to children over various generations yielding an adaptive trajectory. `RNAmutants` considers the entire ensemble of *k* mutants to generate independent samples of stable sequence-structure pairs and thus reveals features such as complex structures that are hard to reach by local methods such as `mateRNAl`.

Our first algorithm `RNAmutants` (see Sec. 4.2) enumerates all mutant sequences and samples the ones with the <u>globally</u> lowest folding energy [71]. It enables us to identify the most stable structures accessible through mutation processes. Noticeably, `RNAmutants` sorts the sequences by the number of mutations separating them from the initial population. We note that `RNAmutants` does not constitute a model of replicating populations but rather a sampling method to obtain representative statistics on the space of stable mutant sequences.

An important element of this work is the interpretation of `RNAmutants` curves where the x-axis shows the number of mutations (e.g. Fig. 3). When this number of mutations is null or very low, the data represents the values we expect from uniformly sampled sequences. By contrast, large numbers of mutations (50 being the maximum number of mutations on sequences of length 50 since it is possible that all positions contain a mutation) are associated with an increased preference for mutants with stable folds and a loss of sequence specificity. The largest hamming neighbourhoods (See Fig. S7) are therefore the ones with strongest thermodynamical pressure.

**Figure 3:**
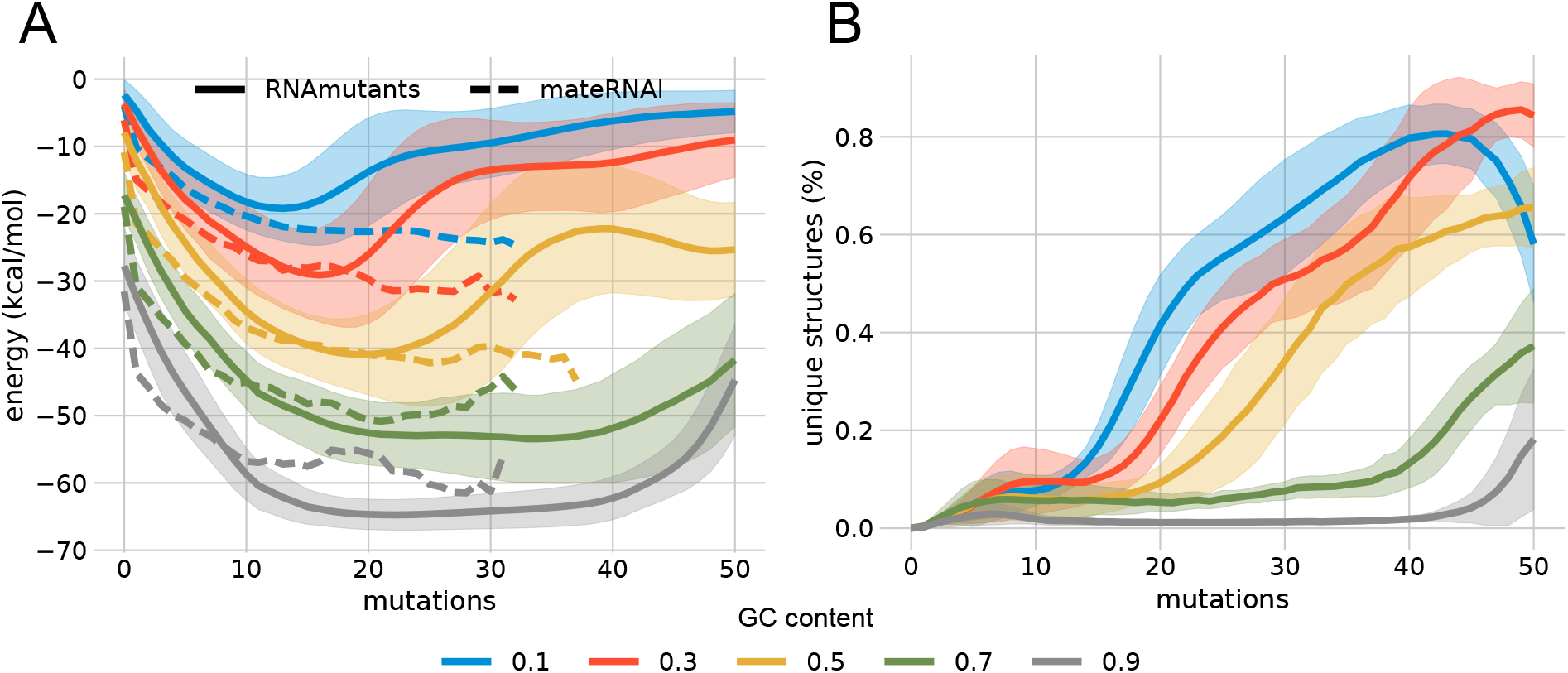
(A) Energy of `RNAmutants` and `mateRNAl` mutational landscape. Shaded regions include one standard deviation of `RNAmutants` energy per mutational distance. Dashed lines mark mean values for `mateRNAl` energies binned by mutations from starting sequence using mutation rate *µ* = 0.02. (B) Fraction of unique structures at every *k* neighbourhood found by `RNAmutants`.

Equipped with a global view of the mutational landscape from RNAmutants, we implemented replication-based algorithms to understand how such a landscape could be traversed by populations. We present `mateRNAl` which has been developed for this study. `mateRNAl` simulates the evolution of a population of replicating RNA sequences that preferentially selects the most stable structures under GC content bias. We also developed variants of this algorithm enabling us to study the antagonistic effect of a negative selection pressure against the most stable structures. Since initial pools of random 50nt sequences are unlikely to contain self-replicating RNA, we frame our experiments in the context of non-catalytic replication [36, 64] which could have been at play until structures with self-replicating abilities are discovered [48].

We study the evolutionary landscape of RNA sequences of length 50 preserving GC contents varying between 0.1 and 0.9 (1 being full G or C sequences). The size of molecules is of particular interest as 50 nucleotides appears to be the upper limit for non-enzymatic self-replicating processes [22], but also because multi-branched structures occur only on RNAs with sizes above 40 (See Fig. 1). It also turns out to the be minimal known size for a natural ribozyme [13]. Shorter ribozymes have been synthesized [66] yet these would not feature complex secondary structure motifs characteristic of most ribozymes.

### 2.2 Energy landscape of RNA mutational networks

We start by characterizing the distribution of folding energies of stable structures accessible from random seeds in the mutational landscape using `RNAmutants`. Our simulations show that, initially, increasing mutational distances from random seeds results in more stable structures at all GC content regimes (See solid lines in Fig. 3). We observe that the folding energies of the samples represent at least 80% of the global minimum energy attainable over all *k* mutations within less than 10 mutations from the seed (see Fig. S3). This suggests that over short evolutionary periods, mutations can play an important stabilizing role as stable structures are likely to be found near the seed.

By contrast, at larger mutational distances (i.e. 15 mutations and above) we notice an increase in the ensemble energy for all GC content regimes. This behaviour is likely due to the exponential growth in sequence space that accompanies higher mutational distances, which results in a more uniform sampling of the ensemble. In this case, less stable sequences outnumber lower energy sequences with high Boltzmann weights (See Fig. S2).

As expected, we observe that the nucleotide content is an important factor in determining the stability of achieved sequence-structure pairs across mutational networks. The accessible sampled energies are strongly constrained by the allowed GC content of sequences in all GC content regimes, whereby higher GC contents favor the sampling of more stable states. However, despite these constraints, all mutational ensembles effectively produce more stable states than the initial state.

Next, we characterize the diversity of structures sampled by RNAmutants. First, we calculate the number of unique structures found in each mutant neighbourhood. Fig. 3B shows two distinct regimes. At low to intermediate GC contents (*≤* 0.5), the immediate neighbourhood (*≤* 10 mutations) of seed sequences is dominated by few stable structures, after what the percentage of unique structures suddenly increases. This sudden change of regime accompanies the increase of the average folding energies observed in Fig. 3A. By contrast, at higher GC contents (*≥* 0.7) the structural diversity increases only at very large mutational distances (*≥* 40), when the footprint from the seed is almost fully lost.

We also explore the repertoire of secondary structure elements represented in the low energy mutational neighbourhoods. Fig. S5 and 4A show the percentage of secondary structures with an internal loop or multi-loop. Although these structural elements are relatively frequent in random sequences (i.e. seeds), they are also not very stable (Fig. 3A). When the mutational distance increases, mutations tend to create stable stem-loop structures and erase irregularities in the original phenotypes. Although, after this stabilization phase, the fraction of internal and multi-branched loops rises again explaining the structural diversity discussed above.

In this work, we focus on the occurrence of multi-branched secondary structures because this motif is often found in functional RNAs such as the hammerhead ribozyme. While previous studies of short randomly sampled RNA sequences (*<* 35 nt.) have shown that simple hairpin structures dominate the low-energy structural landscape, we find that for longer molecules (i.e. 50 nt.) this landscape is enriched with complex multi-branched structures. This finding is in good qualitative agreement with databases of evolved structures where multi-loop structures emerge in families slightly under 50 nucleotides long (Fig. 1A).

Across all runs, we sampled a total of 9419 sequences containing a multi-loop (no more than 1 multi-loop per structure was ever observed as is to be expected for such length scales). Interestingly, we find that unlike internal loops, multi-loops occur under very specific conditions in our sampling. We identify a clear surge in multi-loop frequency at a mutational distance of *∼* 35 (See Fig. 4A), with a mean GC content of *∼* 0.45 (See Fig. 4C). Furthermore, their energy distribution is tightly centered around *∼ −*15 kcal mol^*−*1^ (See Fig. 4B). These values are remarkably close to those of multi-branched structures of similar lengths observed in the Rfam database (See Fig. 1A). In particular, the latter shows a clear bias toward medium GC contents as we identified 148 Rfam families with multi-loops with a GC content of 0.5 (among all Rfam families with sequences having at most 200 nucleotides), but only 80 with a GC content 0.3 and 40 with a GC content of 0.7. This serves as further evidence that GC content is an important determinant of the evolution of structural complexity. It also appears that these features are a general property of the distribution of multi-loops in the mutational landscape given the sequence entropy of the set of sequences containing multi-loops is quite high (0.945 out of 1). This indicates that the observed properties are likely a feature of the GC content bias and not due to over-representation by isolated groups of similar sequences. Further analysis carried out in Section 2.3 suggests that this enrichment of complex structures is not simply an artifact of larger Hamming neighbourhoods that accompanies deeper mutational explorations.

**Figure 4:**
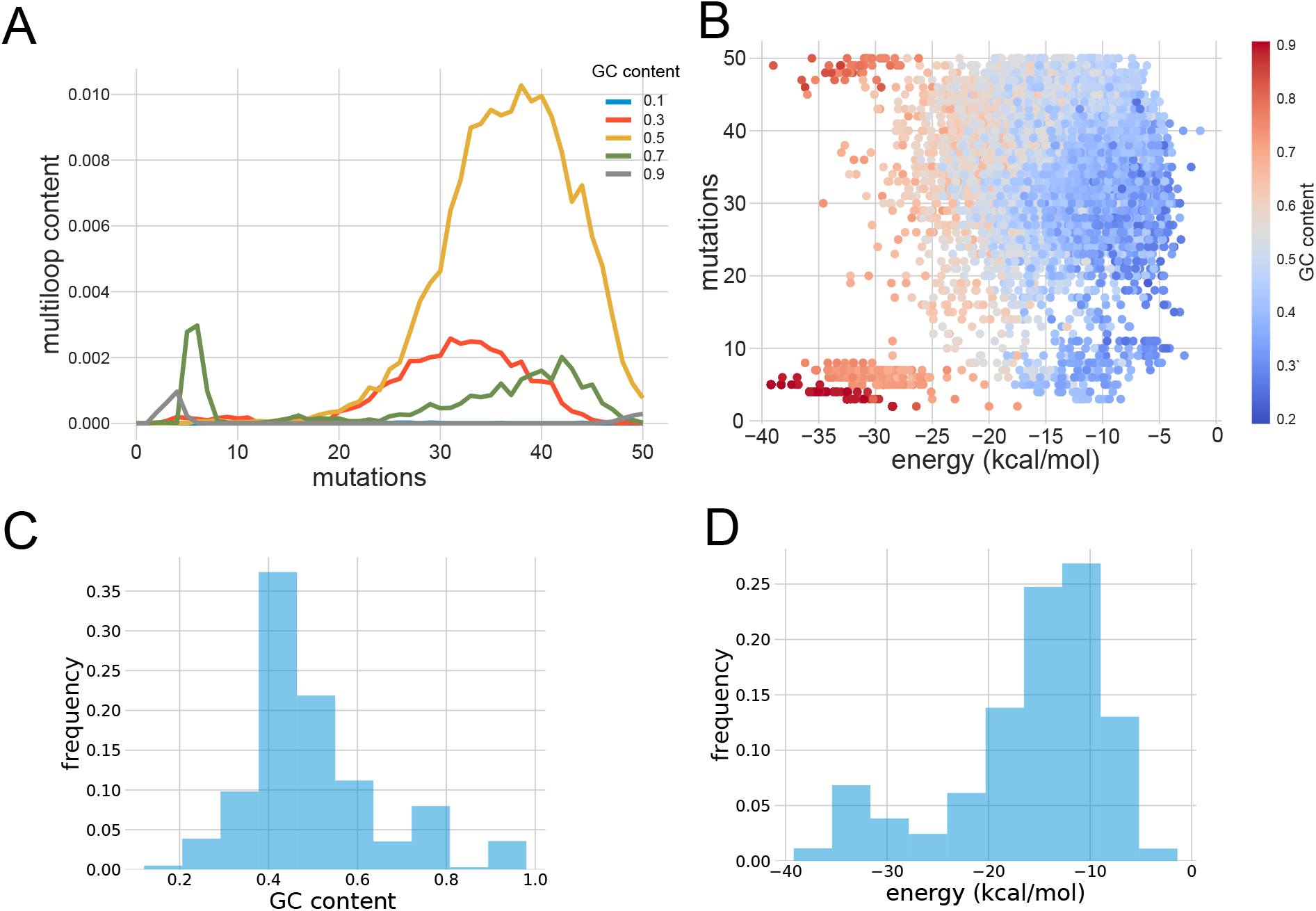
Analysis of the distribution of multi-branched structures in the RNA mutational landscape sampled by `RNAmutants`. (a) The frequency of multi-branched structures with respect to the number of mutations from the seed sequence. (b) Plot of folding energies and GC contents of each individual multi-branched structure. (c) Distribution of the GC content of multi-branched structures. (d) Distribution of folding energies of multi-branched structures.

We also note a smaller peak of multi-loop occurences closer to the seed sequences (*∼* 6 mutations) for higher GC contents around 0.7. Interestingly, with folding energies ranging from *−*25 to *−*40 kcal mol^*−*1^, these multi-branched structures are significantly more stable than those present in the main peak (See Fig. 4B). This is also in agreement with the energies observed in the Rfam database for structures within this range of GC content values (See Fig.1B).

##### Observation 1

- Low and intermediate GC contents (≤ 0.5) promote structural diversity.
- Internal and multi-branched loops are relatively frequent in the low-energy landscape.
- The frequency of stable multi-branched loops in the low-energy landscape is in good qualitative agreement with the Rfam distribution.

### 2.3 Random replication *without* selection

In the previous section, we observed that the low-energy structural landscape is enriched with multi-loop architectures at specific GC contents. We must next determine under which conditions a replicating population can reach this reservoir. Our first scenario aims to study the behaviour and outcome of random replication process <u>without natural selection</u>. Here, we consider a simple model in which RNA molecules are duplicated with a small error rate, but preserving the GC content [65]. In our simulations, we use an error rate of 0.02 (probability of introducing a point mutation) to allow immediate comparison of the number of elapsed generations, and identical transitions and transversions rates. Under these assumptions, we can directly compute the *expected* number of mutations in sequences at the *i^th^* generation (see Methods). Fig. S6 shows the results of this calculation for GC content biases varying from 0.1 to 0.9. Noticeably, our data reveals that after a short initialization phase (i.e. after *∼* 50 generations), sequences with a GC content of 0.5 have on average slightly more than 35 mutations. This observation is in good adequacy with the peak of multi-branched structures identified in Fig. 4A and Fig. 4C. This combination of events suggests that the population is randomly exploring the sequence landscape and gets fixed once stable structures are discovered. Indeed, the mutation distance where multi-loops are observed coincides with the largest mutational neighbourhood (See Fig. S7), thus where the sequence specificity pressure is minimal. We conclude that a simple undirected replication mechanism could explain an enrichment of RNA populations with stable multi-branched structures.

To complete this analysis and assess the different structural compositions between the uniform and low-energy landscapes, we sample sequences at each mutational neighbourhood uniformly at random and compute their minimum free energy (MFE) value and secondary structure. We compare in Fig. 5 the average MFE and frequency of multi-loops in MFE structures between the uniform (“Random”) samples and sequences sampled from the `RNAmutants` low-energy ensemble. Importantly, we report separately the statistics for sequences with or without multi-loops.

**Figure 5:**
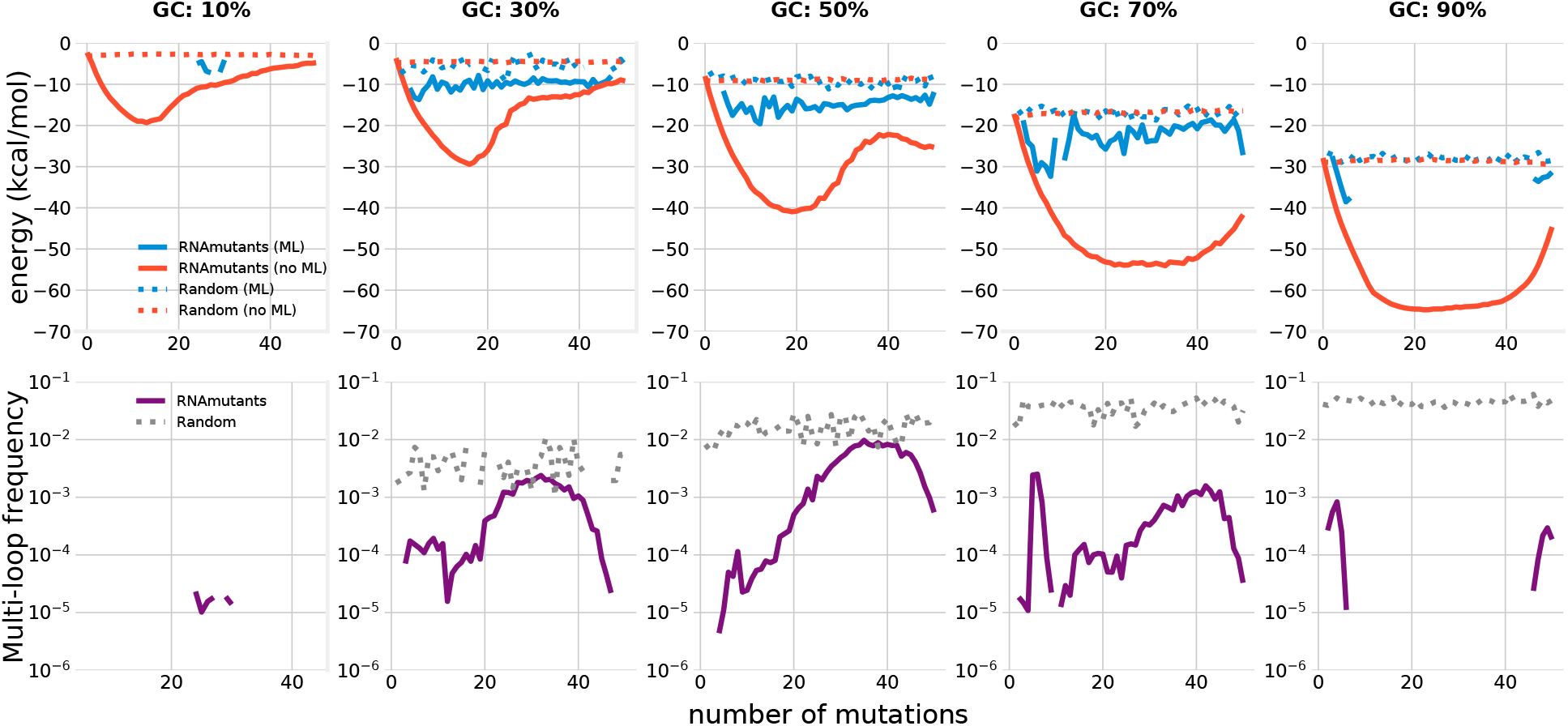
Analysis of multi-loop distributions in uniform and low energy populations. First row: average minimum free energies of uniformly sampled mutants (“Random”; dotted lines) and sequences from the low energy ensemble (“RNAmutants”; plain lines). Sequences with a minimum free energy structure having a multi-loop (“ML”; blue) are separated from the others (“no ML”; orange). Second row: frequency of multi-branched structures in the minimum free energy structure of uniformly sampled mutants (“Random”; dotted grey lines) and sequences sampled from the low energy ensemble (“RNAmutants”; purple lines)

Unsurprisingly, the accumulation of mutations does not impact the results in random populations. First, we note that although multi-branched structures are relatively frequent in random populations (on average between 1 and 10%), they also have higher folding energies than sequences in the low-energy landscape. Higher GC contents tend to increase these trends. This data supports previous observations made on shorter sequences showing that multi-branched structures are rare and relatively unstable in random pools of sequences [60].

By contrast, `RNAmutants` samples exhibit a different pattern. While multi-branched structures are rare in random populations (i.e. no mutations), their frequency increases with the number of mutations (i.e. increased preference toward stable structures). Interestingly, the frequency of stable multi-branched structures almost matches those obtained with random sequences at GC contents between 0.3 and 0.5 and mutational distances from 30 and 40 (second row in the third and fourth columns of Fig. 5).

The analysis of folding energies reveals another interesting phenomenon. While the average energies of multi-branched structures remains steady at all mutational distances, this is not the case of other structures with simpler architectures (first row of Fig. 5). Lower GC content regimes from 0.3 to 0.5 are characterized by a clear increase of average energies at mutational distances over 20 (i.e. MFE structures are less stable), which is not observed at higher GC contents. We conclude from these observations that the relative weight of multi-branched structures in the low energy ensemble (i.e. `RNAmutants`) increases due to a better (collective) resilience of this architecture to point-wise mutations and/or a more uniform distribution in the sequence landscape. In turn, it increases their density in the large/distant mutational neighbourhoods. Moreover, it is worth noting that the values of the folding energies at these GC regimes are also close to those observed in Rfam.

Eventually, we also distinguish a secondary peak of occurrences of multi-branched structures in the vicinity of the seeds (i.e. 5-10 mutations) at higher GC regimes (0.7). By contrast, this higher density appears to result from mutants folding with marginally lower energies. It suggests the presence of mutants with improved fitness to the structures of the seeds rather than a global enrichment of multi-branched structures in these neighbourhoods. We discuss this phenomenon in the next sub-section.

##### Observation 2

- Multi-branched structures are relatively common but unstable in random populations.
- At intermediate GC contents (0.3 − 0.5), multi-branched loops are frequent among the stablest structures.
- The folding energies of stable multi-loop is similar to those observed in databases.

### 2.4 Random replication with selection for stable structures

Our `RNAmutants` and random replication simulations suggest that mutational networks contain reservoirs of complex structures accessible by undirected replication mechanisms. At this point, our main question is to determine if a natural selection process, independent of a particular target, could help reaching these regions.

To address this question we build an evolutionary algorithm named `mateRNAl`, where the fitness is proportional to the folding energies of the molecules. Intuitively, `mateRNAl` simulates the behaviour of a population of RNAs selecting at each generation the most stable sequences regardless of the structure adopted to carry functions (See **Methods 4.1**). This setting is similar to the energy-based selection described by Fontana et al. [15] on binary sequences. These selected structures are therefore by-products of intrinsic adaptive forces.

We start all simulations from random populations of size 1000 and sequences of length 50, and performed 50 independent simulations for each GC content and varying mutation rates. Importantly, we also vary the strength of the selective pressure applied on the population (i.e. the *β* in Eq. 1). All simulations were run for 1000 generations.

We show the frequency and folding energy of multi-branched structures observed during our simulations with `mateRNAl` in Fig. 6. Here, a higher selective pressure (i.e. larger values of *β*) and GC content reduces the frequency of multi-loops but increases the folding energy. Interestingly, a transition occurs when the value of *β* shift from 0.01 to 0.05. By contrast, higher GC contents increase both the frequency of multi-loops and folding energies. Yet, variations of the selective pressure result in more variance of the folding energy.

**Figure 6:**
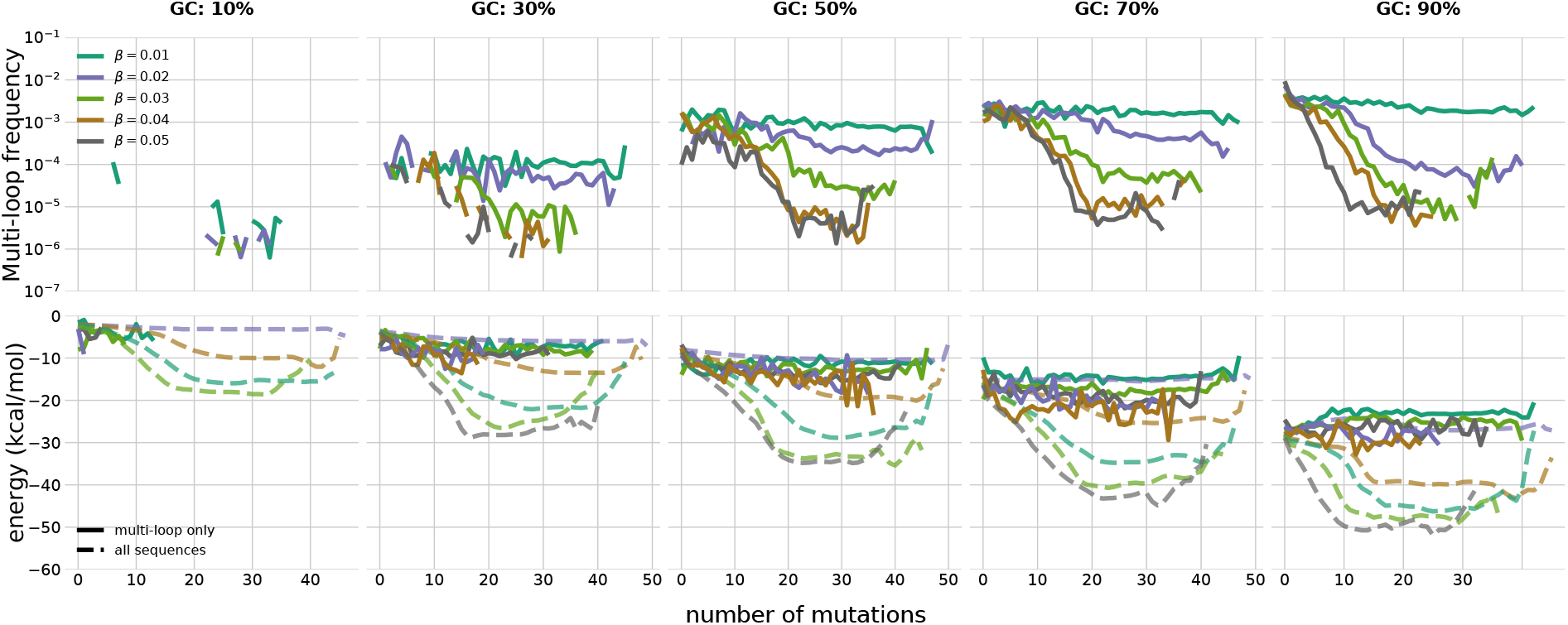
Analysis of multi-loop frequency and energy in mateRNAl populations as a function of mutational distance. First row: average frequency of multiloop structures in populations at each mutational distance. Each line represents a simulation at varying selection strengths (parameter *β*). Subplot columns correspond to simulations at various GC content biases. Second row: Average population energy at each mutational distance. Plain lines report statistics calculated for multi-branched structures only, while dashed lines show the measurements obtained for all structures.

In our simulations, only high GC contents (*≥* 0.7) produce populations with multi-loop frequencies comparable to those observed in databases. It follows that the `mateRNAl` evolutionary scenario does not seem fit to explain how to reach multi-branched sequences with low to intermediate GC content (0.3 *−* 0.5). However, as noted earlier, there is a second pool of multi-branched structures at higher GC content (*≥* 0.7) that are characterized by their proximity to the seed (See Fig.4B, S1) and lower folding energies (See Fig.4D). Under our assumption, the `mateRNAl` scenario appears to be a legitimate candidate to explain how these structures are reached.

We find that `mateRNAl` populations are able to quickly find low energy multi-branched structures (less *∼* 12 mutations from the initial populations; See Fig. 6), with higher GC content regimes leading to more stable structures. Multi-branched structures are mostly found near the seeds (*≤* 20 mutations) and rather in earlier generations.

We also note an interesting dichotomy between populations at 0.7 and 0.9 GC ratios. While the frequency of multi-loops reached by populations at 0.7 roughly matches the one computed by `RNAmutants` (See Fig. 4A), it is not the case for higher GC contents. Indeed, the `mateRNAl` simulations find a significant number of multi-loops not sampled by `RNAmutants`. The folding energies of these structure appears to be similar to the ones found in Rfam (See Fig. 1B).

All together, these observations suggests that a selective pressure on stable structures might help finding stable multi-branched architectures at the highest GC contents (i.e. 0.9) that are otherwise eclipsed by the vast number of stable single stem structures. At 0.7 though this selection mechanism could still be used to accelerate and promote the discovery of these complex structures uniformly distributed in the sequence landscape.

##### Observation 3

- Multi-loops emerge only in populations with highest GC contents (≥ 0.7).
- Highest GC contents help to quickly reach multi-loops in the vicinity of the seeds.
- A selection pressure toward stable structure promote the discovery of rare stable multibranched structures at 0.9.

## 3 Discussion

We provided evidence that in the absence of selective pressure, the structure of the mutational landscape could have helped to promote the emergence of complex RNA phenotypes. To support our hypothesis, we built a comprehensive representation of the mutational landscape of RNA molecules, and investigated scenarios based on distinct hypotheses.

Our results support parsimonious evolutionary scenarios based on undirected molecular replications with occasional mutations. In these simple models, the GC content appears as a key feature in determining the probability of discovering stable multi-branched secondary structures. Our study reveals two distinct phenomena. At low to intermediate GC contents (0.3 *−* 0.5) the distribution of multi-loops in a replicating population eliciting sequences with the most stable structures resembles the one observed in RNA databases. This observation suggests that an evolutionary scenario in which sequences are replicated without selection but where the most stable structures are selected upon their discovery is sufficient to explain the structural complexity needed for the emergence of life at the molecular level.

By contrast, at higher GC contents (*≥* 0.7) the presence of multi-branched structures appears to require the help of a selective pressure to reach these complex phenotypes. In this work, we simulate replicating RNA populations selecting sequences with the most stable structures at each generation. Then, we show that this mechanism enables a quick discovery of multi-loops in the vicinity of random sequences at GC content regimes at or above 0.7. This finding is in agreement with previous theoretical studies that showed that neutral networks percolate the whole sequence landscape [16, 54]. It also suggests a different origin for multi-branched structures at high GC content. In such scenario, a population of molecules would progressively improve the functional efficiency of the molecules by improving their stability.

The preservation of intermediate GC content values appeared to us as a reasonable assumption, which could reflect the availability of various nucleotides in the prebiotic milieu. This nucleotide composition bias can be interpreted as an intrinsic force that favoured the emergence of life. It also offers novel insights into fundamental properties of the genetic alphabet [17]. Incidentally, these observations suggest further investigations into the role of more complex nucleotide distributions [29].

It is worth noting that our scenario remain compatible with further selection mechanisms that may come into play once a functional and stable architecture is identified to rapidly improve active sites [11, 26]. Eventually, our results could also be used to put in perspective earlier findings suggesting that natural selection is not required to explain pattern composition in rRNAs [58].

Our analysis completes recent studies that aimed to characterize fundamental properties of genotype-phenotype maps [19, 32], and showed that their structure may contribute to the emergence of functional molecules [11]. Whereas previous studies focused on characterizing the static genotype-phenotype map of random sequences, we show that the landscape of stable mutants arising from random seeds favours the discovery of complex structures. It also emphasizes the relevance of theoretical models based on a thermodynamical view of prebiotic evolution [38].

The size of the RNA sequences considered in this study has been fixed at 50 nucleotides. This length appears to be the current upper limit for non-enzymatic synthesis [23], and therefore maximizes the expressivity of our evolutionary scenario. Variations of the sizes of populations or lengths of RNA sequences resulting from indels could be eventually considered with the implementation of dedicated algorithms [70]. Although, if these variations remain modest, we do not expect any major impact on our conclusions.

The error rates considered in this study were chosen to match values used in previous related works (e.g [33]). This choice is also corroborated by recent experiments suggesting that early life scenarios could sustain high error rates [44]. Nevertheless, lower mutation rates would only increase the number of generations needed to reach the asymptotic behaviour (See Fig. S6), and thus would not affect our results.

Finally, we emphasize that our results do not exclude the use of more advanced evolutionary mechanisms [7, 57, 62, 63]. Instead, they provide additional evidence supporting an RNA-based scenario for the origin of life, and can serve as a solid basis for further investigations of more sophisticated models.

## 4 Materials and Methods

### 4.1 Evolutionary Algorithm (mateRNAl)

Here we describe an evolutionary algorithm (EA) for energy-based selection with GC content bias. The algorithm is implemented in Python and is freely available at http://jwgitlab.cs.mcgill.ca/cgoliver/maternal.git.

We first define a population at a generation *t* as a set *P*_*t*_ of sequence-structure pairs. We denote a sequence-structure pair as (*ω, s*) such that *s* is the minimum free energy structure on sequence *ω* as computed by the software RNAfold version 2.1.9 [31]. Each sequence is formed as a string from the alphabet 𝔹:= *{A, U, C, G}*. For all experiments we work with constant population size of *|P_t_|* = 1000 *∀t*, and constant sequence length len(*s*_*i*_) = 50 *∀s_i_ ∈ P_t_*. We then apply principles of natural selection under Wright-Fisher sampling to iteratively obtain *P*_*t*+1_ from *P*_*t*_ for the desired number of generations in the simulation

#### 4.1.1 Initial Population

Sequences in the initial population, i.e. generation *t* = 0, are generated by sampling sequences of the appropriate length uniformly at random from the alphabet 𝔹.

#### 4.1.2 Fitness Function

In order to obtain subsequent generations, we iterate through *P*_*t*_ and sample 1000 sequences with replacement according to their relative fitness in the population. Selected sequences generate one offspring that is added to the next generation’s population *P*_*t*+1_. Because we are sampling with replacement, higher fitness sequences on average contribute more offspring than lower fitness sequences. The relative fitness, or reproduction probability of a sequence *ω* is defined as the probability *F* (*ω, s*) that *ω* will undergo replication and contribute one offspring to generation *t* + 1. In previous studies, *F* (*ω_i_, s_i_*) has been typically defined as a function of the base pair distance between the MFE structure of *ω* and a given target structure. However, in our model, this function is proportional to the free energy of the sequence-structure pair, *E*(*ω, s*) as computed by `RNAfold`.

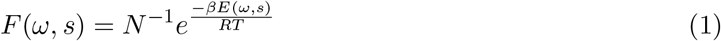

The exponential term is also known as the Boltzmann weight of a sequence-structure pair. *N* is a normalization factor obtained by summing over all other sequence-structure pairs in the population as 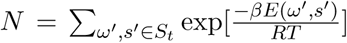. This normalization enforces that reproduction probability of a sequence-structure pair is weighted against the Boltzmann weight of the entire population. *β* is the selection strength coefficient. Higher values of *β* result in stronger selection for stability and vice versa. *R* = 0.000 198 71 kcal mol^*−*1^ K^*−*1^ and *T* = 310.15*K* are the Boltzmann constant and standard temperature respectively.

When a sequence is selected for replication, the child sequence is formed by copying each nucleotide of the parent RNA with an error rate of *µ* known as the mutation rate. *µ* defines the probability of incorrectly copying a nucleotide and instead randomly sampling one of the other 3 bases in 𝔹.

#### 4.1.3 Controlling population GC content

There are two obstacles to maintaining evolving populations within the desired GC content range of *±*0.1. First, an initial population of random sequences sampled uniformly from the full alphabet naturally tends converge to a GC content of 0.5. To avoid this, we sample from the alphabet with probability of sampling GC and AU equal to the desired GC content. This way our initial population has the desired nucleotide distribution. Second, when running the simulation, random mutations are able to move replicating sequences outside of the desired range, especially at extremes of mutation rate and GC content. To avoid this drift, at the selection stage, we do not select mutations that would take the sequence outside of this range. Instead, if a mutation takes a replicating sequence outside the GC range, we simply repeat the mutation process on the sequence until the child sequence has the appropriate GC content. Given that populations are initialized in the appropriate GC range, we are likely to find valid mutants relatively quickly and always avoid drifting away from the target GC.

### 4.2 RNAmutants

The evolutionary algorithm implemented in mateRNAl is similar to a local search. At every time step new sequences are close to the previous population and in particular to the elements with higher <u>fitnes</u>s.

In contrast, `RNAmutants` [71] can sample sequence-structure pairs (*ω, s*) such that (1) the sequence is a *k*-mutant from a given seed *ω*_0_—for any *k*—and (2) the probability of seeing the pair is proportional to its <u>fitness</u> compared to all pairs (*ω*′, *s*^′^) where *w*′is also an *k*-mutant of *ω*_0_.

In addition `RNAmutants` provides an unbiased control of the samples GC content allowing direct comparisons with `mateRNAl`.

We note that although the structure sampled is not in general the MFE replacing them by it does not significantly change the results, as shown in Fig. S2. Therefore we replace the sampled structure with the MFE to simplify the study.

For each GC content in {0.1, 0.3, 0.5, 0.7, 0.9}(*±*0.1) we generated 20 random seeds of length 50. For each seed, at each mutational distance (i.e. number of mutations from the seed) from 0 to 50, at least 10 000 sequence-structure pairs within the target GC content of the seed were sampled from the Boltzmann distribution. The software was run on Dual Intel Westmere EP Xeon X5650 (6-core, 2.66 GHz, 12MB Cache, 95W) on the Guillimin High Performance Computing Cluster of Calcul Québec. It took over 12 000 CPU hours to complete the sampling.

#### 4.2.1 Sequence-structure pairs weighted sampling

Given a seed sequence *ω*_0_, a fixed number of mutations *k*, and the ensemble 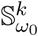 of all pairs sequence-structure such that the sequences are at hamming *k* from *ω*_0_. Similarly to Sec. 4.1.2 the probability of sampling a sequence-structure pair 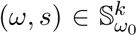 will be its Boltzmann weight, a function of its energy. Formally, if the energy of the sequence *ω* in conformation *s* is *E*(*ω, s*) then the weight of the pair is:

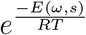

where as before the Boltzmann constant *R* equals 0.000 198 71 kcal mol^*−*1^ K^*−*1^and the temperature *T* is set at 310.15*K*. The normalization factor, or partition function, 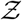 can now be defined as:

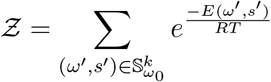

and thus the probability of sampling a pair (*ω, s*) is:

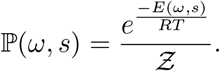

By increasing *k* from 1 to *|ω*_0_*|*(= 50) an exploration of whole mutational landscape of *ω*_0_ is performed. To compute 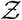 for each value of *k*, `RNAmutants` has a complexity of 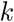. This has to be done only once per seed. The weighted sampling of the sequences themselves has complexity of 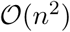.

#### 4.2.2 Controlling sample GC content

Due to the deep correlation between the GC content of the sequence and its energy, the GC base pair being the most energetic in the Turner model [67] which is used by `RNAmutants`, sampling from any ensemble S will be highly biased towards sequences with high GC content. To get a sample (*ω, s*) at a specific target GC content, a natural approach is to continuously sample and reject any sequence not fitting the requirements. Such an approach can yield an exponential time so a technique developed in [72] is applied.

An unbiased sampling of pairs (*ω, s*) for any given GC target can be obtained by modifying the Boltzmann weights of any element (*ω, s*) with a term **w**^*ω*^ ∈ [0, 1] which depends on the GC content of *ω*. At its simplest, it can be the proportion of GC in *ω*. The weight of (*ω, s*) becomes

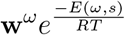

which implies that a new partition function 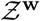 needs to be defined as follows:

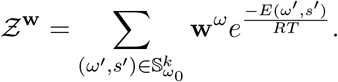

To find the weights **w** for any target GC an exact solution could be found. In practice, the weight **w**^*ω*^ is determined as follows, applying the bissection algorithm. Given the number of GC in *ω*, *|ω|*_GC_(resp number of AU *|ω|*_AU_) and two weight **w**_GC_ and **w**_AU_. We define **w**^*ω*^ as 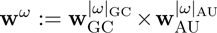.

At the first iteration, a thousand sequences are sampled with **w**_GC_ and **w**_AU_ both as 0.5. If the average GC content on the sampled sequences is too high, the value of **w**_GC_ is decreased by half and **w**_AU_ increased accordingly. If the average GC content is too low, the opposite is done. Each time, the sequences with the desired GC content are kept. This process is repeated until the desired amount of sequences with the required GC content is produced. In practice, after a few iterations almost all sampled sequences contain the target GC content and as shown this sampling is unbiased [72]. An interesting observation is that the same method can be applied to preferentially sample sequences with any other desired feature.

### 4.3 Sequence divergence in a random replication model

We estimate the expected number of mutations in randomly replicated sequences (section 2.3) using the transition matrix defined by K. Tamura [65]. We use a mutation rate *α* = 0.02 mirroring the mutation rate used in `mateRNAl`, and assume that transition and transversion rates are identical. The target GC content is represented with the variable *θ* = {0.1, 0.3, 0.5, 0.7, 0.9}. The transition matrix is shown below.

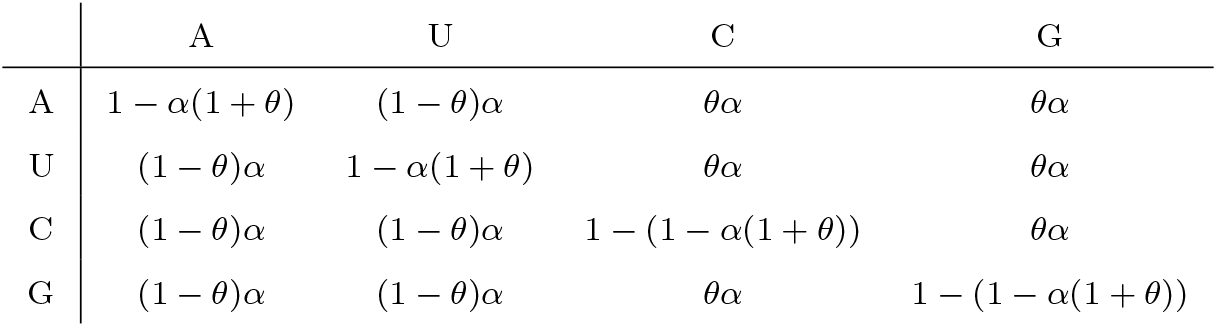

This matrix gives us the transition rate from one generation to the next one. To obtain the mutation probabilities at the *k^th^* generation, we calculate the *k^th^* exponent of this matrix. Then, we sum the values along the main diagonal to estimate the probability of a nucleotide to be the same at the initial and *k^th^* generation.

## 5 Author contributions

CGO, VR, and JW designed the research, analyzed the results, and wrote the manuscript. CO and VR conducted the computational experiments.

## Supplemental Materials

## 1 Sampling schemes

`RNAmutants` samples sequence-structure pairs according to their ensemble weight. This method therefore does not guarantee the sampling of the minimum free energy (MFE) structure for a given sequence. However, for a given “suboptimal” sampling we can use `RNAfold` to obtain the MFE structure for every sequence, producing a set of sequence-structure pairs that can be directly compared to `mateRNAl` data which samples only MFE structures. In Fig. S2 we show that this procedure produces sequence-structure pairs with nearly identical ensemble frequencies, allowing us to proceed with MFE structures in downstream analysis.

## 2 Energy landscapes

With free energy being the driving force in our simulations, we next compared mean energy values for mutation populations in `mateRNAl` and `RNAmutants`. We show in Fig. S3 that a random starting seed in `RNAmutants` is on average under 15 mutations away from the global minimum of its mutational ensemble. This feature of the energy of mutational landscapes is exploited by the local search nature of `mateRNAl` which is able to rapidly identify stable solutions with very few mutations. More specifically, we tested various mutation rates (*µ*) for mateRNAl and superimposed mean values to those obtained by `RNAmutants`, Fig. S4. We note that higher mutation rates lead to deeper mutational explorations yet never reaching as far as `RNAmutants`. Regardless, it appears that the most stable structures are maintained at lower mutation rates, suggesting a tradeoff between exploration and refinement.

## 3 Structural complexity

Here we summarize the structural features discovered by mateRNAl and RNAmutants. In this study we distinguish between three major structural features: stack, internal loop, and multiloop. The stack is produced from consecutive base pairing interactions and thus higher stack numbers form longer stem structures. These represent the simplest of the structural motifs. The internal loop occurs when a stacking is interrupted by unpaired bases forming a loop like structure within a stem. A multi-loop is also an unpaired region that forms a loop but which connects three or more stems, also known as a junction. Fig. S5 shows the effect of GC content on the occurence of internal looping structures as a function of mutational distance in `RNAmutants` simulations.

## 4 Analysis of Hamming neighbourhoods

We provide complementary data to characterize the sequences available at specific mutational distances and GC contents. Fig. S6 shows the number of mutations accumulated in populations evolving under a random model preserving the GC content. We use for this purpose the model introduced by K. Tamura [65] (See Sec. 4.3). Fig. S7 indicates the number of sequences with exactly *k* mutations and with a GC content within *±*0.1 of the target GC content. Thus, it shows the size of the neighbourhoods.

---

**Figure S1:**
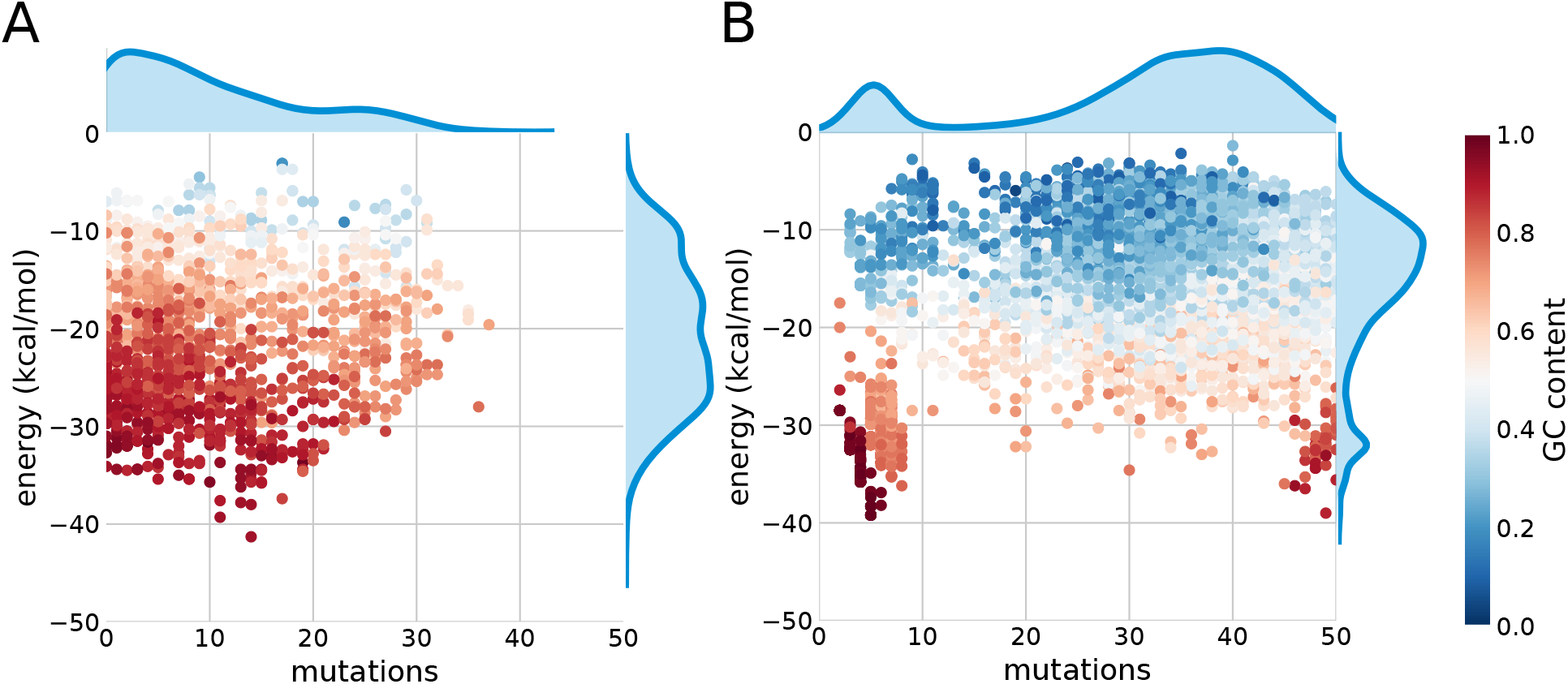
Distribution of multi-loop occurrences for `mateRNAl` (A) with selection pressure *β* = 0.04 and `RNAmutants` (B).

**Figure S2:**
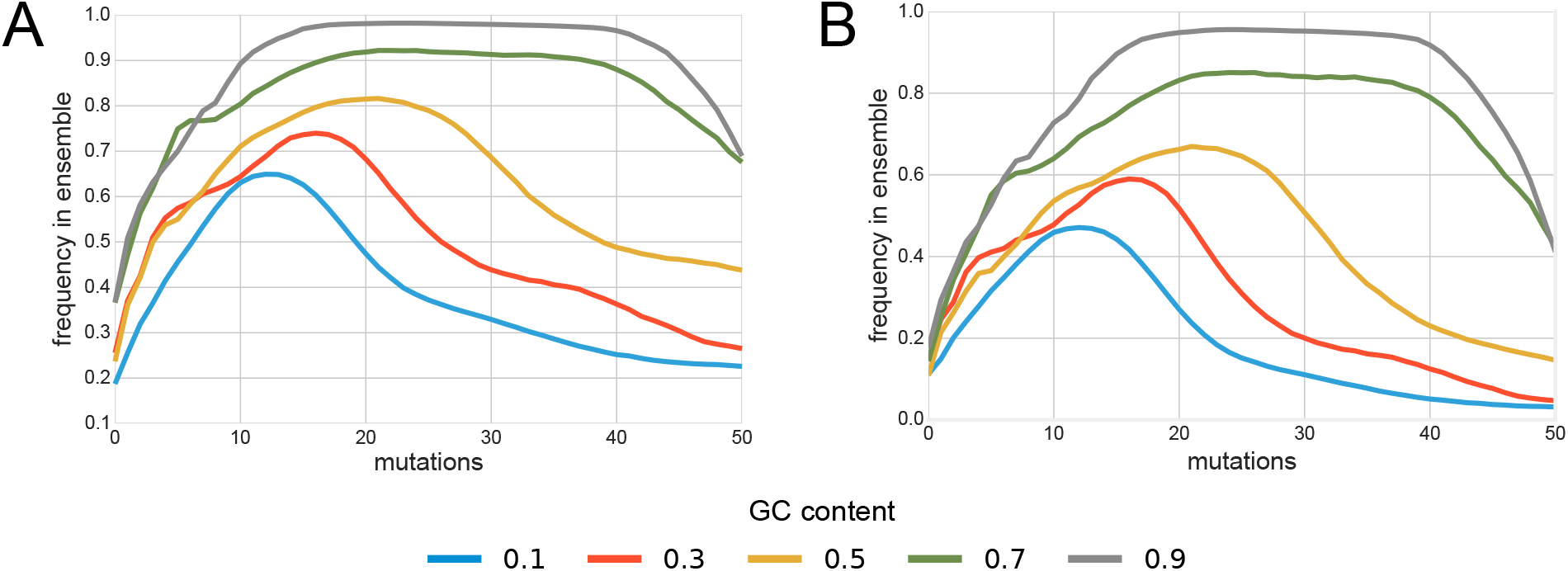
Frequency of sequence-structure pair in Boltzmann ensemble for MFE (A) and suboptimal sampling (B) in `RNAmutants`. Since the `RNAmutants` is not guaranteed to sample the MFE structure for all sequences, we compute the MFE structure using RNAfold for all the sampled sequences and plot the resulting energies.

**Figure S3:**
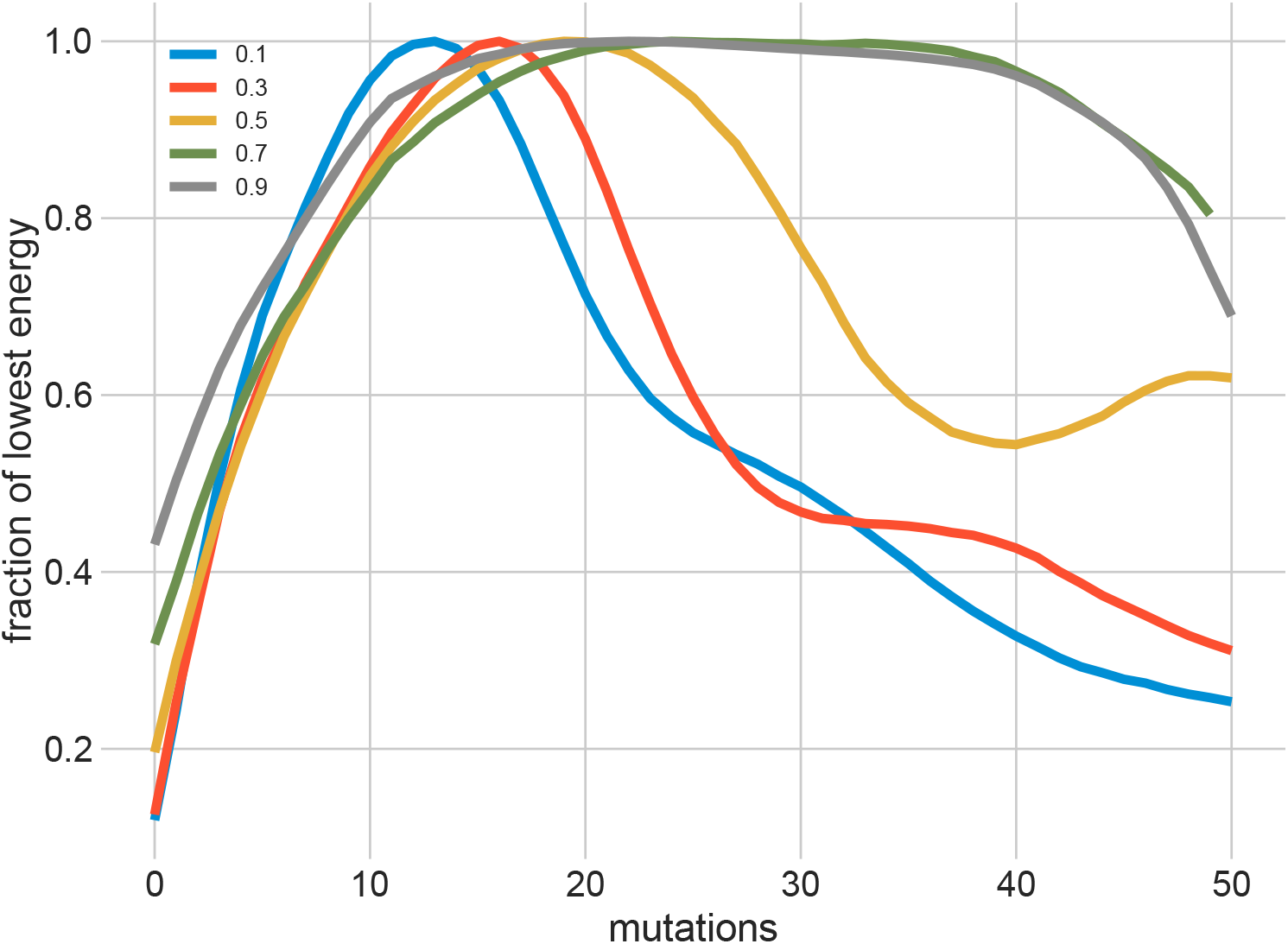
Percentage of the global minimum energy reached by `RNAmutants` by mutational distance.

**Figure S4:**
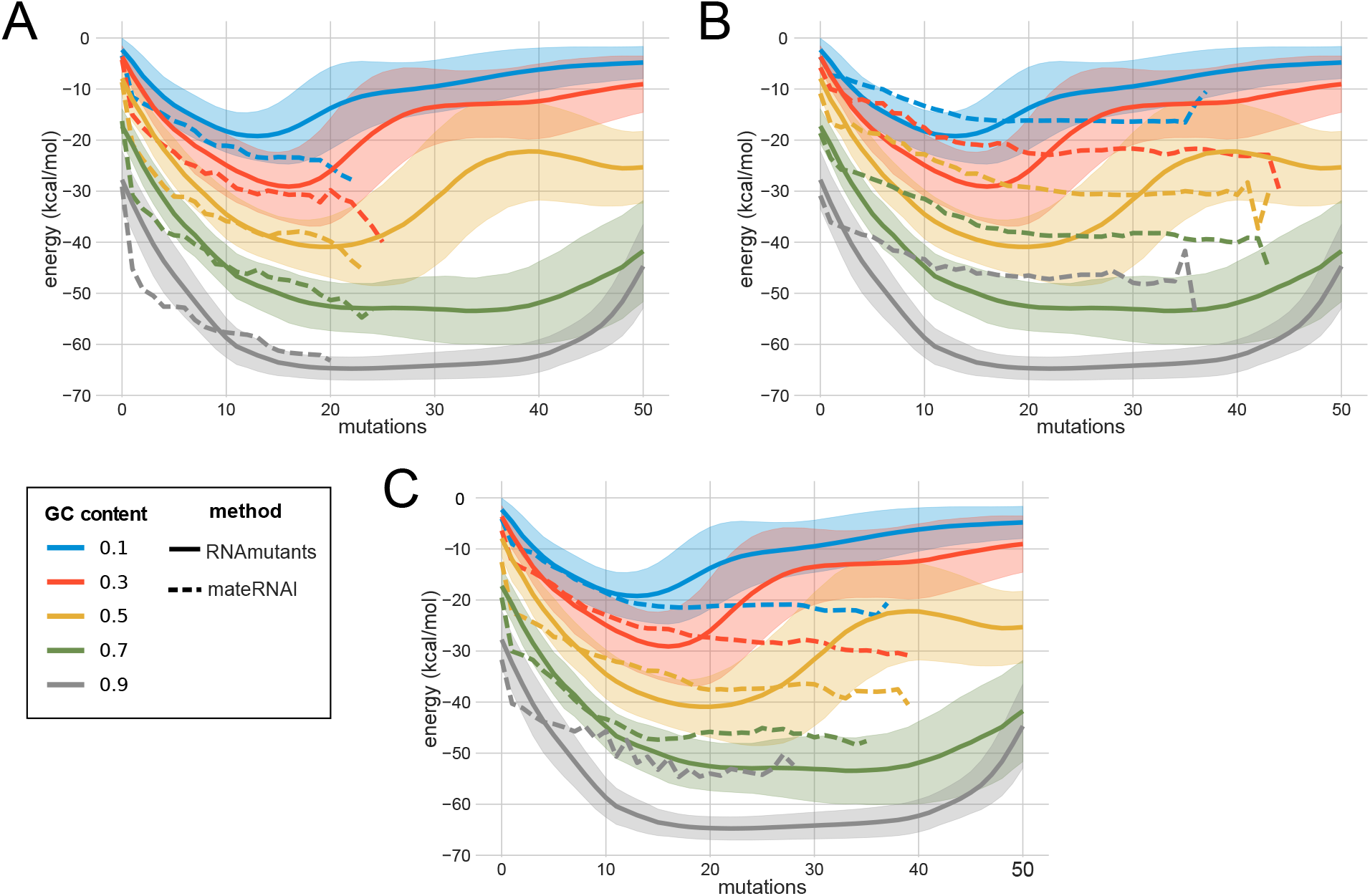
Average energy comparison between `mateRNAl` and `RNAmutants` using different mutation rates for `mateRNAl`.

**Figure S5:**
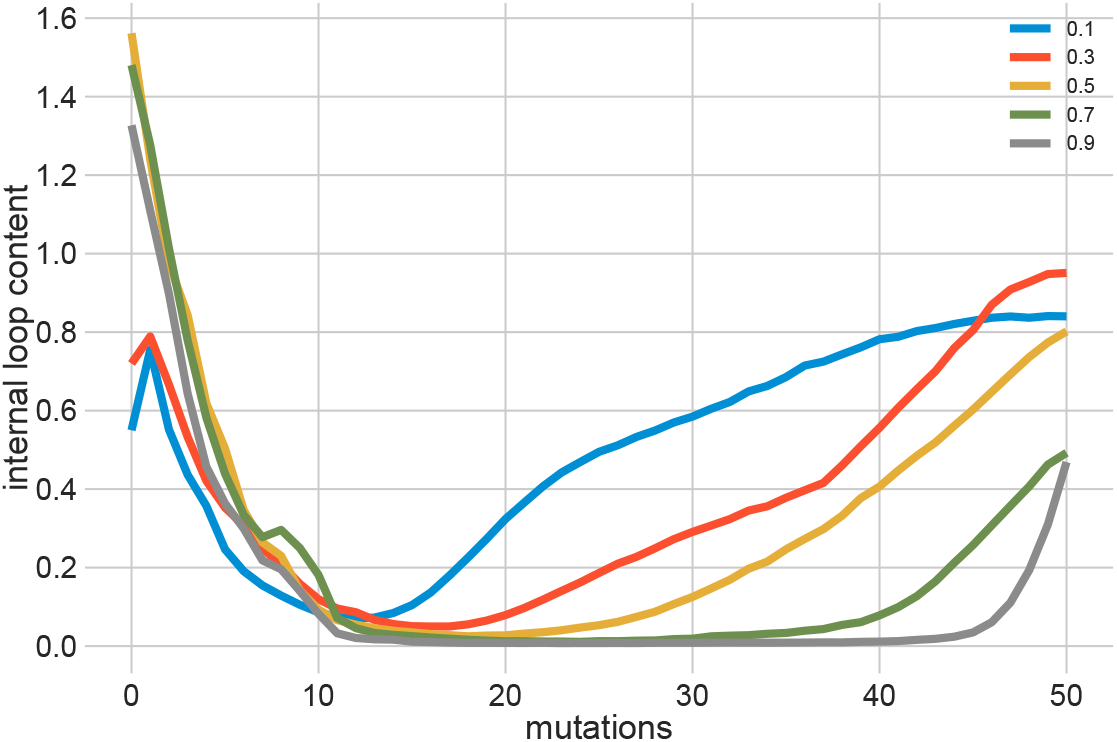
Fraction of *k*-mutant neighbourhoods populated with internal looping structures in `RNAmutants`.

**Figure S6:**
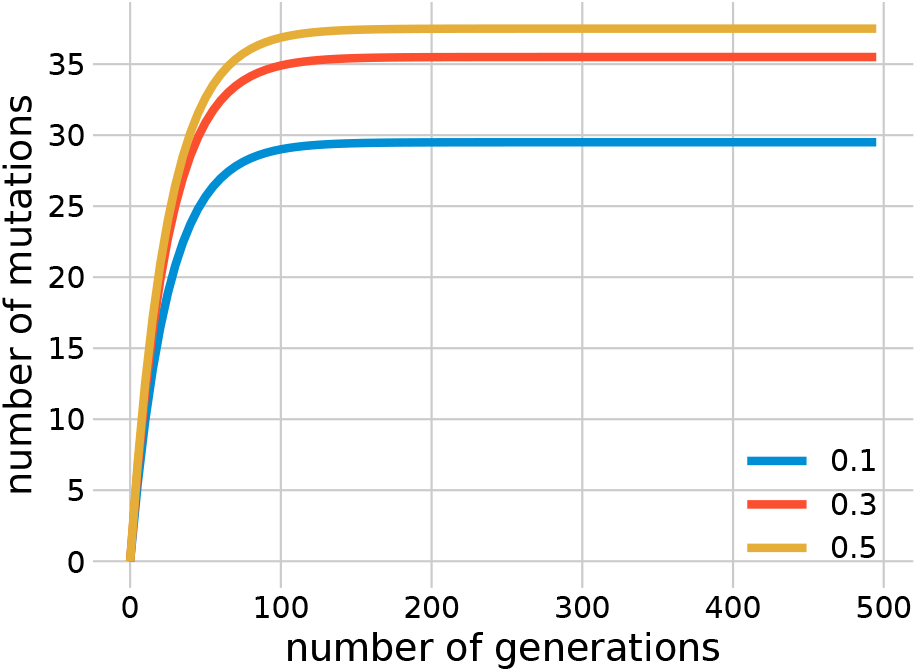
Number of mutations accumulated in populations evolving under a random GC-preserving replication model.

**Figure S7:**
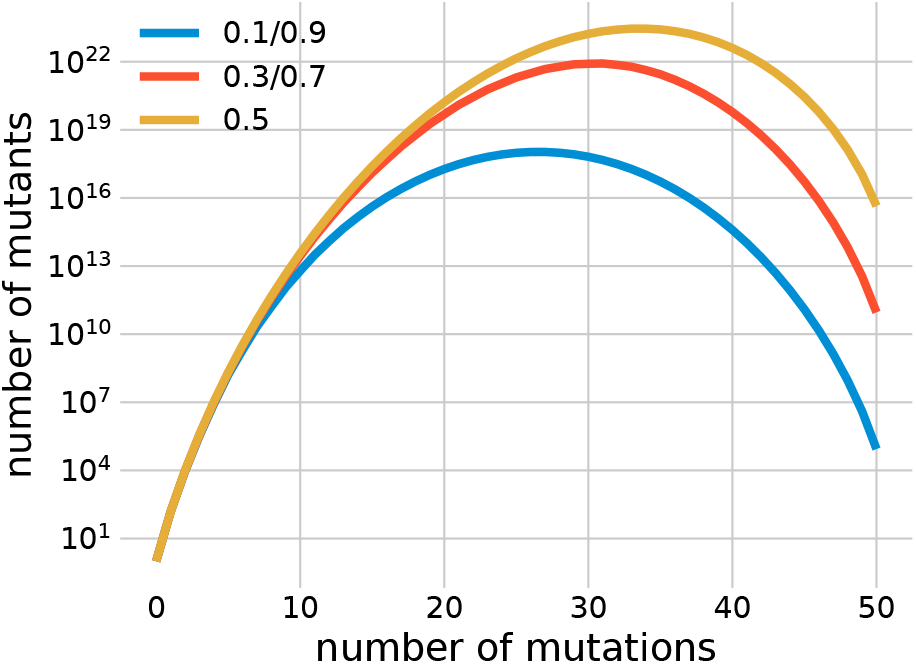
Number of sequences in hamming neighbourhoods.

